# Paleo-Eskimo genetic legacy across North America

**DOI:** 10.1101/203018

**Authors:** Pavel Flegontov, N. Ezgi Altinişik, Piya Changmai, Nadin Rohland, Swapan Mallick, Deborah A. Bolnick, Francesca Candilio, Olga Flegontova, Choongwon Jeong, Thomas K. Harper, Denise Keating, Douglas J. Kennett, Alexander M. Kim, Thiseas C. Lamnidis, Iñigo Olalde, Jennifer Raff, Robert A. Sattler, Pontus Skoglund, Edward J. Vajda, Sergey Vasilyev, Elizaveta Veselovskaya, M. Geoffrey Hayes, Dennis H. O’Rourke, Ron Pinhasi, Johannes Krause, David Reich, Stephan Schiffels

## Abstract

Paleo-Eskimos were the first people to settle vast regions of the American Arctic around 5,000 years ago, and were subsequently joined and largely displaced around 1,000 years ago by ancestors of the present-day Inuit and Yupik. The genetic relationship between Paleo-Eskimos and Native American populations remains uncertain. We analyze ancient and present-day genome-wide data from the Americas and Siberia, including new data from Alaskan Iñupiat and West Siberian populations, and the first genome-wide DNA from ancient Aleutian Islanders, ancient northern Athabaskans, and a 4,250-year-old individual of the Chukotkan Ust'-Belaya culture. Employing new methods based on rare allele and haplotype sharing as well as established methods based on allele frequency correlations, we show that Paleo-Eskimo ancestry is widespread among populations who speak Na-Dene and Eskimo-Aleut languages. Using phylogenetic modelling with allele frequency correlations and rare variation, we present a comprehensive model for the complex peopling of North America.

---

Current evidence suggests that present-day Native Americans descend from at least four distinct streams of ancient migration from Asia^1–3^. The largest ancestral contribution was from populations that separated from the ancestors of present-day East Asian groups ~23,000 calendar years before present (calBP), occupied Beringia for several thousand years, and then moved into North and South America approximately 16,000 calBP^2^. To be consistent with the previous genetic literature we call this lineage “First Americans”, while acknowledging that indigenous scholars have suggested the term “First Peoples” as an alternative. The deepest phylogenetic split in this group gave rise to one lineage that contributed to northern North American groups (including speakers of Na-Dene, Algonquian and Salishan languages), and to another lineage that is found in some North Americans as well as all Native Americans from Mesoamerica southward^1,2,4^. The 12,600 calBP ancient genome from an individual assigned to the Clovis culture belongs to the southern lineage^5^. In addition, a separate source of Asian ancestry that has been called “Population Y” contributed more to Native American groups in Amazonia than to other Native Americans^2,3^. A third stream of migration contributed up to ~50% of the ancestry of the Inuit and Aleut peoples (Eskimo-Aleut speakers), but the Asian source population for this stream remained unidentified^1^. Of key importance for understanding the impact of these different lines of ancestry are populations speaking Na-Dene languages, which include the Tlingit, Eyak (recently extinct), and Northern and Southern Athabaskan languages, spoken across much of Alaska and northwestern Canada, with additional isolated Na-Dene languages spoken further south along the Pacific Coast and in southwestern North America^6^. It has been argued^1^ that Na-Dene-speaking populations harbor ancestry from another distinct migration: ancient Paleo-Eskimos deriving from Chukotka around 5,000 calBP and expanding throughout the American Arctic for more than 4,000 years^7–9^. An alternative view is that Paleo-Eskimo-derived ancestry disappeared entirely from temperate North America after the arrival of Thule Inuit, and the distinctive ancestry in Na-Dene speakers might instead reflect admixture from Thule Inuit^2,8,10^.

The archaeological record in the Arctic provides clear evidence for the spread of Paleo-Eskimo culture, which spread across the Bering strait about 5,000 calBP^9,11–13^, and expanded across coastal Alaska, Arctic Canada and Greenland a few hundred years later. Direct ancient DNA data has proven that the Paleo-Eskimo cultural spread was strongly correlated with the spread of a new people^7,8^ that continuously occupied the American Arctic for more than four millennia until ~700 calBP^9,14,15^. A long-term cultural, and likely linguistic and genetic, boundary was established upon their arrival, which separated populations in the coastal Arctic tundra from indigenous Native American groups who populated the interior forest zone and were plausibly ancestors of present-day Na-Dene speakers^16^. Paleo-Eskimo archeological cultures are grouped under the Arctic Small Tool tradition (ASTt), and include the Denbigh, Choris, Norton, and Ipiutak cultures in Alaska and the Saqqaq, Independence, Pre-Dorset, and Dorset cultures in the Canadian Arctic and Greenland^9^. The ASTt source has been argued to lie in the Syalakh-Bel’kachi-Ymyakhtakh culture sequence of East Siberia, dated to 6,500 - 2,800 calBP^17,18^. In this paper, we use the genetic label “Paleo-Eskimo” to refer to the ancestry associated with ancient DNA from the ASTt and “Neo-Eskimo” to refer to ancient DNA from the later Northern Maritime tradition. While we recognize that some indigenous groups would prefer that the term “Eskimo” not be used, we are not aware of an alternative term that all relevant groups prefer instead. The terms “Paleo-Inuit” and “Thule Inuit” have been proposed as possible replacements for “Paleo-Eskimo” and “Neo-Eskimo”, respectively^19^, but the use of “Inuit” in this context might seem to imply that individuals from these ancient cultures are more closely related to present-day Inuit than to present-day Yupik, whereas genetic data show that Yupik and Inuit derive largely from the same ancestral populations (see below). Moreover, the term “Thule” does not cover the whole spectrum of Northern Maritime cultures, being strongly associated with the latest phase of this tradition. We therefore use the “Eskimo” terminology here while acknowledging its imperfections.

Paleo-Eskimo dominance in the American Arctic ended about 1,350 – 1,150 calBP, when the Thule culture became established in Alaska and rapidly spread eastwards after 750 – 650 calBP^9,14,15^. This spread has been shown genetically to reflect the movement of people^8^. The Thule Inuit had material culture links to hunter-gatherer societies in the Bering Strait region (e.g., Old Bering Sea culture, starting about 2,200 calBP, and Birnik culture), who depended on marine resources^20^. More complex and diverse transportation technologies, weaponry, and, most importantly, a food surplus created by whale hunting, contributed to the success of these Neo-Eskimo cultures and to eventual disappearance of the Paleo-culture with which it competed^11,15,21^.

A 4,000-year-old Paleo-Eskimo from western Greenland, associated with the Saqqaq culture, was the first ancient anatomically modern human to have his whole genome sequenced, yielding a genome of 16x coverage^7^. Later work reported low-coverage data for additional individuals affiliated with the Pre-Dorset, Dorset and Saqqaq cultures^8^. These studies showed that Paleo-Eskimos were a genetically continuous population^8^ and are most closely related, among present-day groups, to Chukotko-Kamchatkan-speaking Chukchi and Koryaks who live in far eastern Siberia^2,7,8^. The split time between the first Saqqaq individual sequenced and the Chukchi was estimated at 6,400 – 4,400 calBP^7^, consistent with archaeological data. Present-day speakers of Eskimo-Aleut languages and ancient Neo-Eskimos represent another continuous population, related to Paleo-Eskimos and Chukchi, but distinct^8^. No admixture was detected between Neo- and Paleo-Eskimos in the Canadian Arctic and Greenland^8^, consistent with the lack of evidence for interactions between their material cultures^14^. However, Raghavan *etal.*^8^ hypothesized early gene flow from the Neo-Eskimo into the Paleo-Eskimo lineage in Beringia, and Raff *etal.*^22^ found mitochondrial evidence for possible gene flow from Paleo-Eskimos into the ancestors of contemporary Iñupiat from the North Slope of Alaska. It is important to recognize that substantial coverage genome-wide data from Alaskan Paleo-Eskimo cultures, including Choris and Norton, and from Chukotkan cultures possibly related to Paleo-Eskimos (the Ust’-Belaya and Wrangel island sites) have never been reported.

In this study, we resolve the debate around the distinctive ancestry in Na-Dene and determine the genetic origin of Neo-Eskimos and their relationships with Paleo-Eskimos and Chukotko-Kamchatkan speakers. We present the first genomic data for ancient Aleutians, ancient Northern Athabaskans, Chukotkan Neo- and Paleo-Eskimos, and present-day Alaskan Iñupiat. We also present new genotyping data for West Siberian populations (Enets, Kets, Nganasans, and Selkups). Analyzing these data in conjunction with an extensive set of public sequencing and genotyping data, we demonstrate that the population history of North America was shaped by two major admixture events between Paleo-Eskimos and the First Americans, which gave rise to both the Neo-Eskimo and Na-Dene populations.

## Results

### Dataset

We generated new genome-wide data from 11 ancient Aleutian Islanders that date from 2,320 to 140 calBP, three ancient Northern Athabaskans (McGrath, Upper Kuskokwim River, Alaska, 790 – 640 calBP), two Neo-Eskimos of the Old Bering Sea culture (Uelen, Chukotka, 1,970 – 830 calBP), and one individual of the Ust’-Belaya culture (Ust’-Belaya, Chukotka, 4,410 – 4,100 calBP) (Table 1, Supplementary Table 1, Supplementary Information sections 1 and 2). For each of these 17 individuals, we extracted bone powder in a dedicated clean room, extracted DNA^23^, and prepared a double-stranded library treated with uracil-DNA glycosylase enzymes to greatly reduce the rate of characteristic ancient DNA damage^24^. We enriched the libraries for a targeted set of approximately 1.24 million single nucleotide polymorphisms (SNPs)^25^. We assessed the authenticity of the samples based on the rate of matching of sequences to the mitochondrial consensus, X chromosome polymorphism in males, and cytosine-to-thymine mismatch to the human reference genome in the terminal nucleotides of each read, which is a characteristic signature of genuine ancient DNA (Table 1, Supplementary Information section 3). By itself, this dataset increases the number of individuals from the American Arctic and from far eastern Siberia with more than 1.0x coverage on analyzed positions by 11-fold (10 samples in our study meet this threshold compared to only one that met this threshold previously^7^). In addition to the newly reported ancient data, we report new SNP genotyping data for present-day populations: 35 Alaskan Iñupiat (Inuit), 3 Enets, 19 Ket, 22 Nganasan, and 14 Selkup (Supplementary Table 2).

**Table 1.** Summary of genome-wide data from 17 newly reported samples

| ID1 | ID2 | Skeletal element | Date, calBP (95% CI) | Label | Location | Country | Latitude | Longitude | Sex | mtDNA | Y haplo-group | Data type | Coverage | SNPs |
| --- | --- | --- | --- | --- | --- | --- | --- | --- | --- | --- | --- | --- | --- | --- |
| I1526 | barrow 8, skeleton 4 | molar | 4410 – 4100 | Ust'-Belaya | Ust'-Belaya II, Chukotka | Russia | 65.48 | 173.29 | M | C4a1a3 | Q1a2a | Capture | 4.523 | 832,452 |
| I1525 | burial 13, inventory no. 172 | molar | 1970 – 1590 | Old Bering Sea | Uelen, Chukotka | Russia | 66.19 | -169.9 | F | A2a |  | Capture | 1.031 | 608,585 |
| I1524 | burial 22, inventory no. 163 | molar | 1180 – 830 |  |  |  | 66.17 | -170.75 | M | A2a | Q1a2a1a1 | Capture | 2.618 | 797,816 |
| I5319 | MT_1 | pars petrosa | 790 – 640 | Athabaskan | Tochak McGrath, Upper Kuskokwim River, Alaska | USA | 62.95 | -155.59 | M | A2a1 | Q1a2a1a1 | Capture | 5.731 | 865,897 |
| I5320 | MT_2 |  | 790 – 640 (from MT_1) |  |  |  |  |  | M | A2+(64) | Q1a2a1a1 | Capture | 5.038 | 839,833 |
| I5321 | MT_3 |  | 790 – 640 (from MT_1) |  |  |  |  |  | F | A2+(64) |  | Capture | 4.851 | 827,885 |
| I0721 | 378628 | bone (rib) | 2320 – 1900 | Paleo-Aleut | Chaluka Midden, Umnak Island, Aleutian Islands | USA | 52.99 | -168.82 | M | D2a1a | CT | Capture | 0.025 | 28,805 |
| I0712 | 378623 |  | 1270 – 930 |  |  |  |  |  | F | D2a1a |  | Capture | 0.431 | 395,958 |
| I1126 | 378622 |  | 1250 – 780 |  |  |  |  |  | M | D2a1a | Q | Capture | 0.092 | 103,481 |
| I0719 | 378620 |  | 770 – 390 |  |  |  |  |  | M | D2a1a | Q1a2a | Capture Shotgun | 3.432<br>2.700 | 927,083<br>290,049 |
| I1125 | 378544 |  | 760 – 490 | Neo-Aleut | Ship Rock Island, Aleutian Islands | USA | 53.37 | -167.83 | M | D2a1a | Q1a2a1a1 | Capture | 1.335 | 640,629 |
| I1127 | 377814 |  | 630 – 310 |  | Kagamil Island Warm Cave, Aleutian Islands | USA | 52.99 | -169.71 | F | D2a1a |  | Capture | 1.354 | 662,276 |
| I1128 | 377915 |  | 610 – 290 |  |  |  |  |  | F | D2a1a |  | Capture | 0.433 | 347,752 |
| I1129 | 377917 |  | 600 – 270 |  |  |  |  |  | M | D2a1a | Q1a2 | Capture | 0.687 | 495,889 |
| I1118 | 377811 |  | 530 – 230 |  |  |  |  |  | F | D2a1a |  | Capture | 3.551 | 759,975 |
| I1123 | 377918 |  | 520 – 140 |  |  |  |  |  | M | A2a | Q1a | Capture | 0.128 | 136,906 |
| I1124 | 377919 |  | 500 – 140 |  |  |  |  |  | M | D2a1a | Q1a2a | Capture | 0.156 | 164,481 |

We merged the newly reported ancient and modern data with previously published data to create three main datasets covering Africa, Europe, Southeast Asia, Siberia, and the Americas (Fig. 1, Supplementary Tables 3, 4). For most analyses, we combined groups into meta-populations, as indicated in Fig. 1 and summarized in Supplementary Table 3. The breakdown of groups into these meta-populations was guided by unsupervised clustering using *ADMIXTURE* (Extended Data Fig. 1), *fineSTRUCTURE* (Extended Data Fig. 2), Principal Component Analysis (PCA) (Fig. 2, Extended Data Fig. 3, Supplementary Information section 4). For naming the Arctic meta-populations, we use names of recognized language families: Na-Dene, Eskimo-Aleut, Chukotko-Kamchatkan. We chose these terms since genetic and linguistic relationship patters are highly congruent in this region (see below).

**Figure 1.**
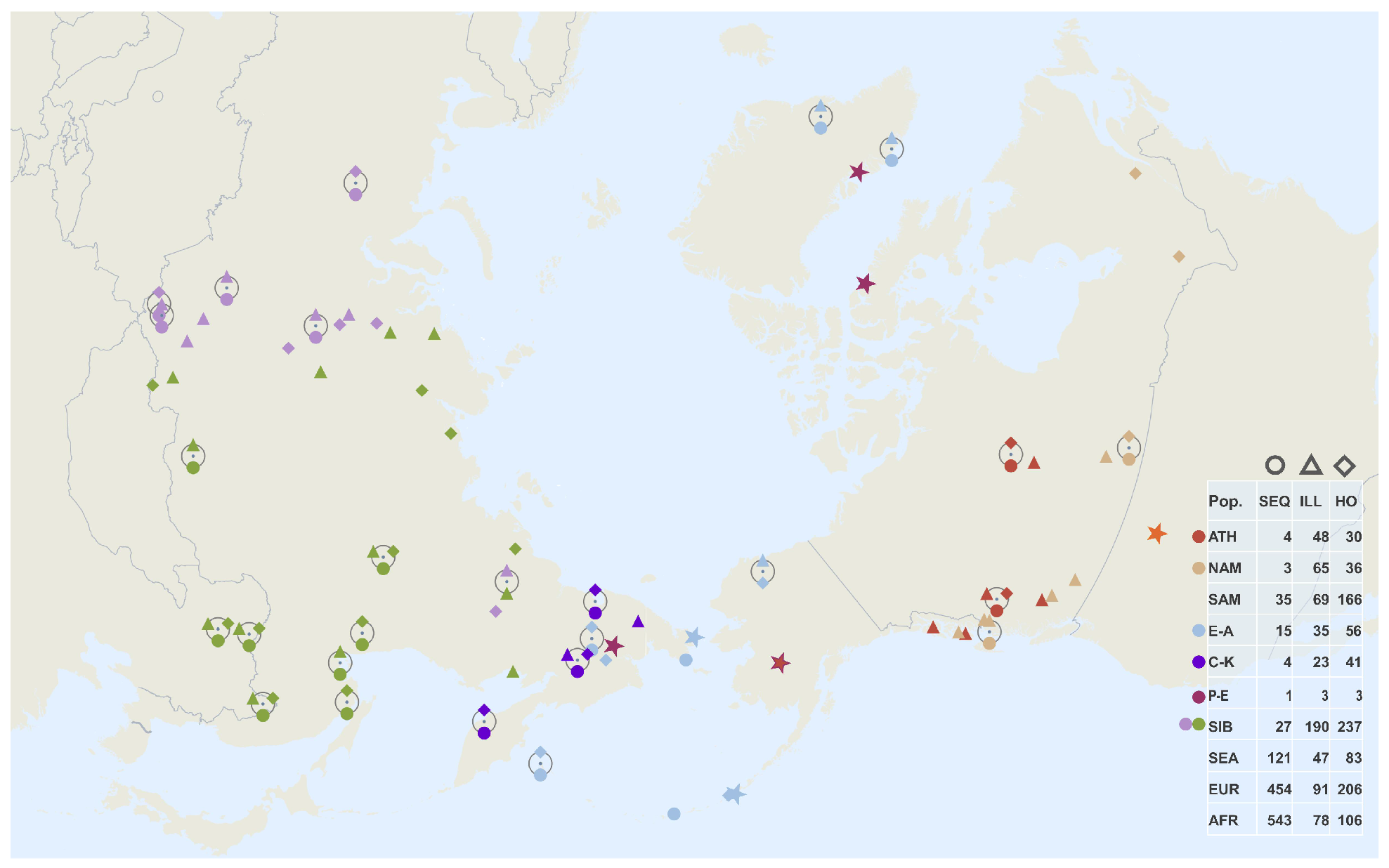
Geographic locations of Siberian and North American populations used in this study. Three main datasets are as follows (Supplementary Tables 3,4): 1) a set based on the Affymetrix Human Origins genotyping array, including diploid genotypes for the ancient Saqqaq^7^ and Clovis^5^ individuals, together with SNP capture data from six ancient Aleuts who had the highest coverage, two unrelated ancient Athabaskans, two ancient Chukotkan Neo-Eskimos, and the Ust’-Belaya Chukotkan Paleo-Eskimo (Table 1); 2) a set based on various Illumina arrays, including Saqqaq and the other ancient samples, and 3) a whole genome data set of 1,207 individuals from 95 populations, including the Clovis, Saqqaq, and one ancient Aleut individual for which we generated a complete genome with 2.7x coverage. The dataset composition, i.e. number of individuals in each meta-population, is shown in the table on the right. Locations of samples with whole genome sequencing data (SEQ) are shown with circles, and those of Illumina (ILL) and HumanOrigins (HO) SNP array samples with triangles and diamonds, respectively. Meta-populations are color-coded in a similar way throughout all figures and designated as follows: Na-Dene speakers (abbreviated as ATH), other northern First Americans (NAM), southern First Americans (SAM), Eskimo-Aleut speakers (E-A), Chukotko-Kamchatkan speakers (C-K), Paleo-Eskimos (P-E), West and East Siberians (WSIB and ESIB), Southeast Asians (SEA), Europeans (EUR), and Africans (AFR). Locations of the Saqqaq, Clovis and other ancient samples are shown with asterisks colored to reflect their meta-population affiliation.

**Figure 2.**
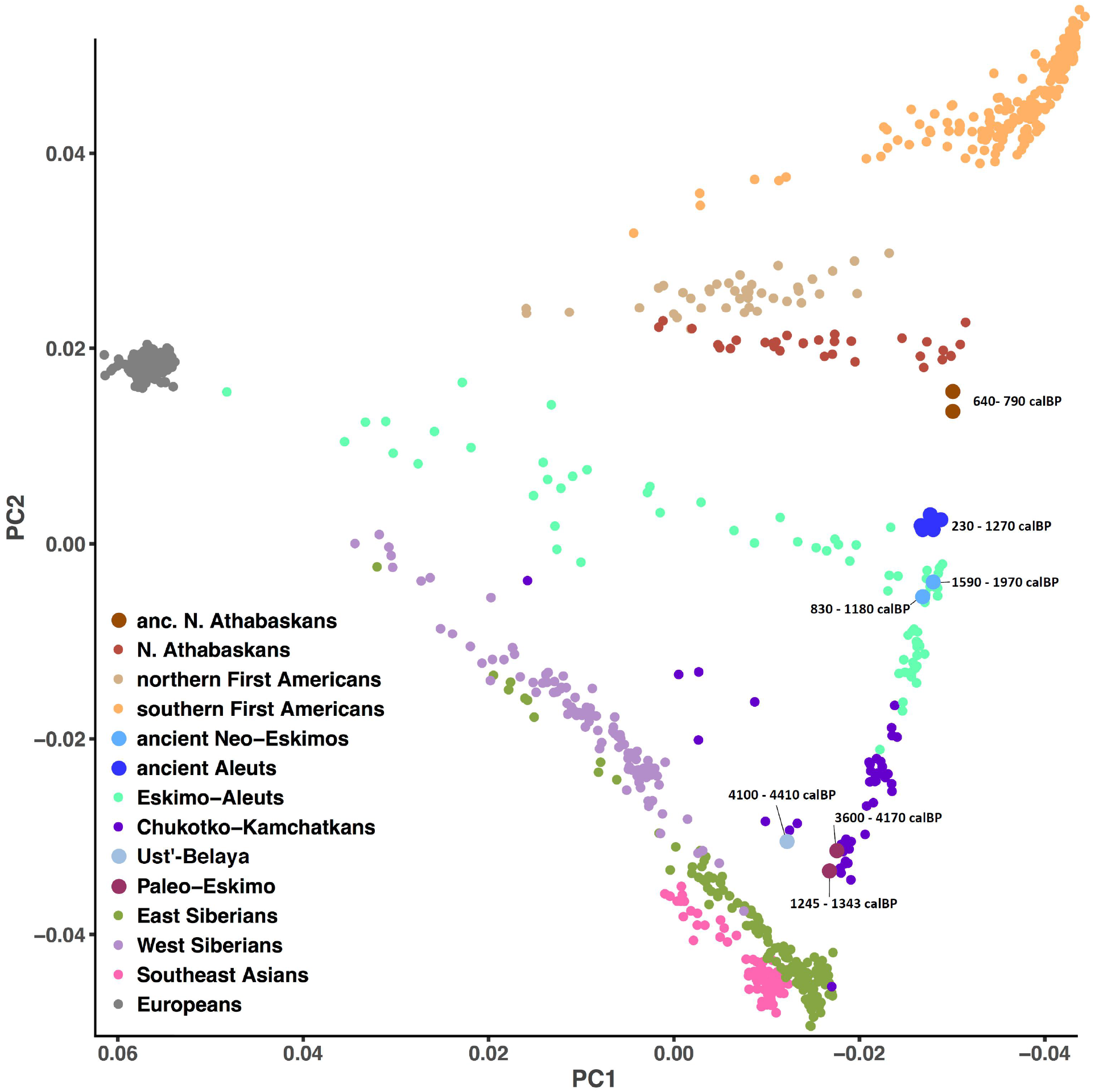
Principal component analysis (PCA) on the HumanOrigins datasets prior to any outlier removal. A plot of two principal components (PC1 vs. PC2) is presented. Calibrated radiocarbon dates in calBP are shown for ancient samples (large circles). For individuals, 95% confidence intervals are shown, and for populations, minimal and maximal dates among all confidence intervals of that population are shown. A similar plot for the Illumina dataset is displayed in Extended Data Fig. 3, and plots for the datasets used for *qpWave/qpAdm* analyses are shown in Supplementary Information section 4. In those datasets, First American, Chukotko-Kamchatkan-speaking and Eskimo-Aleut-speaking individuals having >1% European, African, or Southeast Asian ancestry according to *ADMIXTURE* were removed.

### Gradient of Paleo-Eskimo-related ancestry

PCA applied to SNP array datasets (Fig. 2) reveals a striking linear cline with Paleo-Eskimos (Saqqaq and Late Dorset) and some Chukotko-Kamchatkan speakers at one extreme, then Chukchi, then contemporary Eskimo-Aleut speakers and ancient Neo-Eskimos and Aleuts, then Na-Dene speakers, then northern North Americans, and finally southern First Americans at the other extreme. The patterns were qualitatively identical for the HumanOrigins and Illumina datasets, in analyses carried out with or without transition polymorphisms (Fig. 2, Extended Data Fig. 3, Supplementary Information section 4). This qualitative pattern in PCA is driven by admixture, as we verified using the *qpWave* method^1^. *qpWave* relies on a large matrix of *f4*-statistics measuring allele sharing correlation rates between all possible pairs of a set of outgroups and all possible pairs of a set of test populations. A statistical test^1^ can then be performed to determine whether allele frequencies in the test populations can be explained by one, two, or more streams of ancestry derived in different ways from the outgroups; this test gives a single P-value that appropriately corrects for multiple hypothesis testing. We verified that all the individuals on the PCA cline could be modeled as descended from two streams of ancestry relative to a diverse set of Siberians, Southeast Asians, Europeans, and Africans. Since Chukotko-Kamchatkan speakers are closely related to Paleo-Eskimos as shown here (Fig. 2, Extended Data Figs. 2, 3) and in previous studies^2,7^, we included them along with the American groups as test populations. With this setup, a great majority of all possible population quadruplets of the form (First American, Na-Dene, Eskimo-Aleut, Paleo-Eskimo) were consistent with two streams of ancestry derived from the outgroups (P>0.05), especially on the datasets lacking transition polymorphisms in order to avoid possible confounding effects due to ancient DNA degradation (Supplementary Information section 5).

Under the assumption that the populations at the extremes of the cline are descended solely from one of the source populations, we can assign admixture proportions to all populations in the middle of the cline using *qpAdm*, an extension of *qpWave^26^.* Thus, we attempted modelling diverse American populations as descended from both southern or northern First Americans and Paleo-Eskimos. This analysis reveals a gradient of Paleo-Eskimo-related ancestry proportions, with the relative values almost perfectly proportional to the position along the PCA gradient (Fig. 2, Extended Data Fig. 3). The *qpAdm* estimates of Paleo-Eskimo-related ancestry are as follows: southern First Americans (by definition 0%), northern First Americans (3%), present-day Na-Dene (722%), ancient Northern Athabaskans (23-38%, depending on the dataset), Eskimo-Aleuts other than Yupik (30-68%), Yupik (71-76%), Chukotko-Kamchatkans (~100%), and Paleo-Eskimos (by definition 100%) (Fig. 3, Extended Data Figs. 4, 5). Adding a Chukotko-Kamchatkan-speaking population without recent American back-flow (Koryak) to the outgroup dataset changed these results: three streams of ancestry generally fit the data in the full datasets, but the picture was more ambiguous in the transition-free datasets, with the HumanOrigins-based transition-free dataset still supporting the model with two migration streams (Supplementary Information section 5). Nevertheless, admixture proportions inferred by *qpAdm* remained largely unchanged (Extended Data Figs. 4, 5).

**Figure 3.**
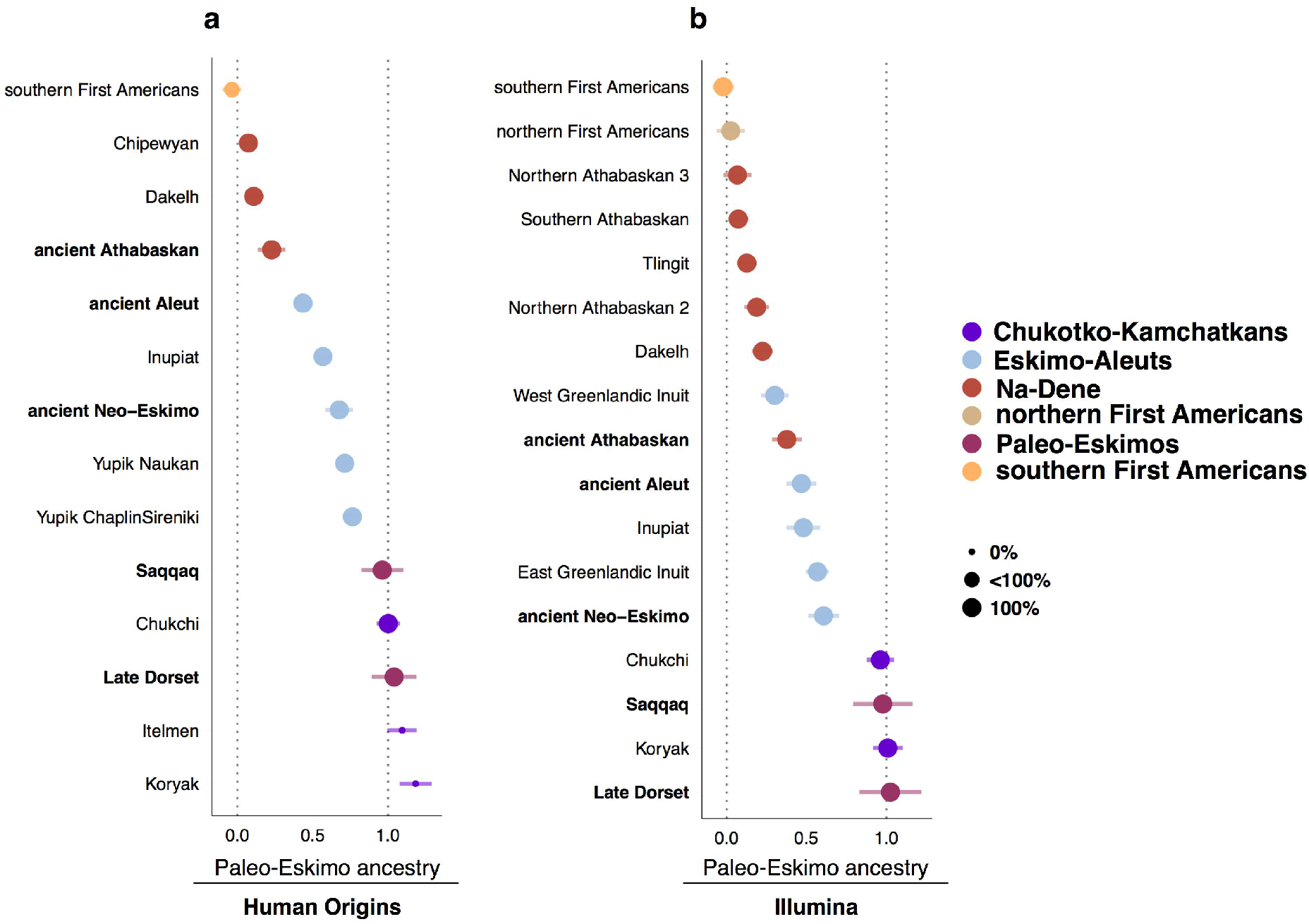
A gradient of Paleo-Eskimo ancestry in America revealed using the *qpAdm* approach. American, Chukotkan, and Kamchatkan populations were modelled as descended from both First American and Paleo-Eskimo sources on the HumanOrigins (**a**) and Illumina (**b**) datasets without transition polymorphisms. First, population triplets were tested with *qpWave* for consistency with two or three streams of ancestry derived from outgroups. Second, *qpAdm* was used to infer admixture proportions in present-day or ancient (in bold) target populations. Saqqaq was considered as a Paleo-Eskimo source for all populations apart from Saqqaq itself, for which Late Dorset was used as a source, and alternative First American sources were selected among the largest populations with little or no detectable admixture: Mixe, Guarani, or Karitiana for the HumanOrigins dataset; Nisga’a, Mixtec, Pima, or Karitiana for the Illumina dataset. Admixture proportions and their standard errors were averaged across triplets including these different First American sources, or across many alternative target populations in the case of southern and northern First Americans. Meta-populations are color-coded according to the legend on the right. Proportion of population triplets consistent with two migration streams is coded by the circle size: small (0%), medium (>0% and <100%), and large (100%). The following sets of outgroups were used: 8 diverse Siberian populations (Nganasan, Tuvinian, Ulchi, Yakut, Even, Ket, Selkup, Tubalar) and a Southeast Asian population (Dai) on the HumanOrigins dataset; 9 Siberian populations (Buryat, Dolgan, Evenk, Nganasan, Tuvinian, Yakut, Altaian, Khakas, Selkup) and Dai on the Illumina dataset. See results for other outgroup sets and for datasets including transitions in Extended Data Figs. 4 and 5.

In summary, all indigenous populations of North America, Chukotka and Kamchatka are consistent with deriving from two ancestry streams to the limits of our resolution, which we term First American and proto-Paleo-Eskimo (PPE). This “distant perspective” treats the region west of the Bering Strait (notably Chukotka and Kamchatka) as part of the American radiation. Usage of a close outgroup within the PPE radiation (Koryak), as also done in Reich *etal.*^1^, yields a “close perspective” and models additional population structure within the PPE radiation which explains the finding in that study of three streams of ancestry connecting Asia to the Americas rather than the two streams of ancestry we focus on here. We find that the PPE source population for Eskimo-Aleut speakers is a distinct line of ancestry, different from Paleo-Eskimos *sensu stricto*, in that it is more closely related to present-day Chukotko-Kamchatkan speakers (see the demographic modelling results below). In contrast, the PPE source that contributed to Na-Dene is most closely related to Paleo-Eskimos *sensu stricto*, as seen by a *qpWave* analysis on population triplets (Na-Dene, First American, Paleo-Eskimo), which are generally consistent with two migration streams on all datasets even with Koryak in the outgroups (Extended Data Figs. 4, 5). Below, we use methods based on autosomal haplotypes and rare variants to further investigate whether Paleo-Eskimos^1^ or Thule Inuit^2,8^ contributed the distinctive ancestry which these analyses show were present in Na-Dene speakers.

### Source of distinct ancestry in Na-Dene

To investigate Paleo-Eskimo ancestry in Native Americans in a hypothesis-free way, we considered haplotypes shared with the ancient Saqqaq individual. As compared to allele frequencies at unlinked loci, autosomal haplotypes in some cases have more power to distinguish potential closely related sources of gene flow^27,28^, such as Thule Inuit and Paleo-Eskimos. Cumulative lengths of shared autosomal haplotypes were produced with *ChromoPainter v.1* for all pairs of individuals^29^. First, for each American individual, we considered the length of haplotypes shared with Saqqaq (in cM), which we refer to as Saqqaq haplotype sharing statistic or HSS. We also estimated haplotype sharing between each American individual and African, European, Siberian, and Arctic (Chukotko-Kamchatkan- and Eskimo-Aleut-speaking) individuals by averaging HSS across members of a given meta-population. To correct for potential biases caused by sequence quality and coverage, the Saqqaq HSS was divided by the African HSS for each group, and the resulting statistic was termed relative HSS (Extended Data Fig. 6).

In both genome-wide genotyping datasets, most Native American individuals with the highest relative Saqqaq HSSs belonged to the Na-Dene group. This enrichment cannot be explained by either Arctic or European admixture in these individuals, as shown by the poor correlation with Arctic and European relative HSSs (Extended Data Fig. 6). We note that some correlation of the Saqqaq and Arctic HSSs is expected under any admixture scenario since Saqqaq falls into the Arctic clade in trees based on haplotype sharing patterns (Extended Data Fig. 2).

While the HumanOrigins dataset includes only two Northern Athabaskan-speaking groups from Canada (Chipewyans and Dakelh) and only three other northern First American groups (Algonquins, Cree, Ojibwa), the Illumina dataset includes six such populations in addition to all extant major branches of the Na-Dene language family: four groups of Northern Athabaskan speakers, one Southern Athabaskan group, and one Tlingit group. At least one individual from each Na-Dene branch demonstrates a relative Saqqaq HSS surpassing that of any Central or South American (Extended Data Fig. 6). The results were very similar when using a genetically distant meta-population (African) and a much closer one (Siberian) as normalizers (Supplementary Information section 6).

To interpret haplotype sharing in a more quantitative way, we analyzed putative admixture events in Na-Dene speakers using *GLOBETROTTER*^30^. To make a complex ancestry history of Na-Dene amenable to *GLOBETROTTER* analysis, we pre-selected individuals based on low European admixture and high Saqqaq HSS (selected individuals are marked in Supplementary Information section 6). Consistent with our qualitative observations, Paleo-Eskimos (represented by the Saqqaq individual) and First Americans were identified by *GLOBETROTTER* as the most likely sources of ancestry for Na-Dene, with the Paleo-Eskimo contribution ranging from 7% to 51%, depending on the dataset and *GLOBETROTTER* setup. Admixture dates were estimated as 2,202 – 479 calBP (Supplementary Information section 7).

As an independent test, we analyzed rare genetic variants in the complete genome dataset. Rare variants, with global frequency of less than 1%, have been shown to have more power to resolve subtle relationships than common variants^31,32^. We calculated rare allele sharing statistics (RASS), which measure the number of rare variants (up to allele frequency 0.2% in the entire dataset) an individual shares with reference meta-populations, in this case Siberian and Arctic (Extended Data Fig. 7, Supplemental Information section 4). To normalize coverage differences between individuals and dataset-specific variant calling biases, we divided these statistics by allele sharing with Europeans or Africans (Extended Data Fig. 7). On a two-dimensional plot combining Arctic and Siberian RASS, four metapopulation lines are visible: Siberian, First American, Chukotko-Kamchatkan, and Eskimo-Aleut (Fig. 4). All four Northern Athabaskan (Dakelh and Chipewyan) individuals are shifted on the Arctic axis by more than three standard error intervals from the First American cluster. The Arctic/Siberian RASS ratios are almost identical in Athabaskans and Saqqaq, but significantly different in present-day Eskimo-Aleut-speaking individuals and in an ancient Aleut individual, for which we generated whole genome shotgun data of 2.7x coverage (Table 1, Fig. 4). Allele sharing statistics behave linearly under recent admixture, and we used linear combinations to calculate expected statistics for First American/Saqqaq and First American/Eskimo-Aleut admixture. Notably, relative RASSs for both Dakelh individuals match those of the simulated First American/Saqqaq admixture, but the statistics for two Chipewyans are consistent with both admixture scenarios (Extended Data Fig. 8). Similar results were obtained in an analysis without transitions (Extended Data Fig. 8).

**Figure 4.**
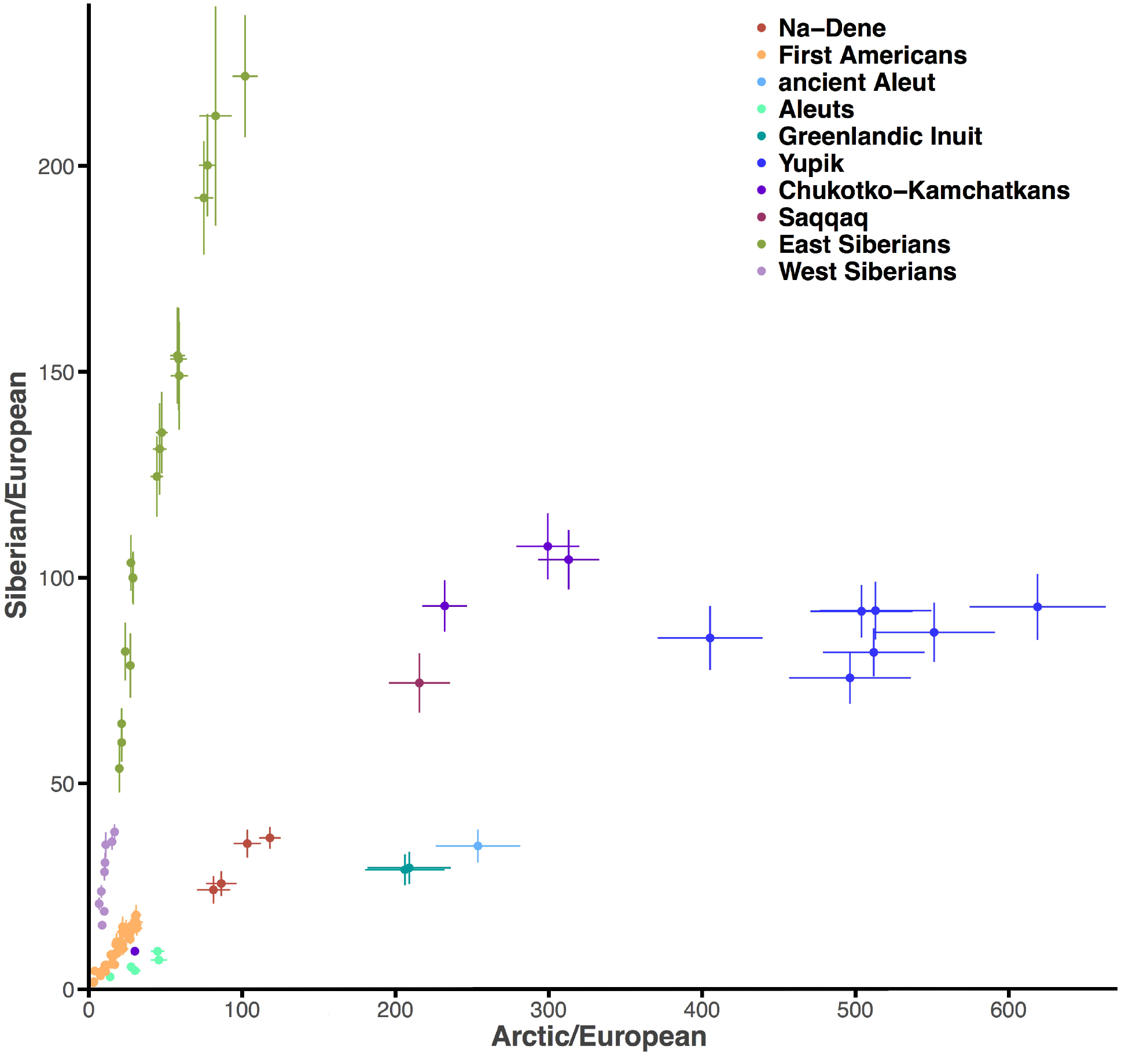
Relative rare allele sharing statistics calculated for each present-day or ancient American individual and the Arctic and Siberian meta-populations. Non-reference alleles occurring from 2 to 5 times in the dataset of 1,207 diploid genomes contributed to the statistics. To take care of variability in genome coverage across populations and of dataset-specific SNP calling biases, we normalized the counts of alleles shared by a given American individual and the Arctic or Siberian meta-populations by similar counts of alleles shared with Europeans. Standard deviations were calculated using a jackknife approach with chromosomes used as resampling blocks, and single standard error intervals are plotted. Plots on the dataset without transitions are shown in Extended Data Fig. 8c,d.

Taken together, our results from several analyses show remarkable consistency: PCA, haplotype and rare allele sharing, *GLOBEROTTER*, and *qpWave/qpAdm* suggest that present-day Na-Dene speakers lacking post-Columbian admixture have roughly 10% to 25% Paleo-Eskimo ancestry. Our newly reported data from the three ancient individuals from the Tochak McGrath site dated at ~800 calBP, found in a region currently inhabited by Na-Dene speakers, are derived from the same combination of First American and Paleo-Eskimo lineages as present-day Na-Dene, providing support for the hypothesis of local population continuity, also supported by continuity in material culture^16^. However, the ancient Tochak McGrath samples have a higher estimated proportion of Paleo-Eskimo ancestry than any present-day Na-Dene speakers in our dataset, 25-40%, suggesting that ongoing gene flow from neighboring First American populations has been reducing the Paleo-Eskimo ancestry in Na-Dene. Paleo-Eskimo ancestry is likely present at a low level in other northern First Americans (Extended Data Figs. 4-6) due to this bidirectional gene flow. Two Dakelh individuals with genome sequencing data available^8^ have yielded consistent results throughout all analyses (Figs. 2-4, Extended Data Figs. 4-8, Supplementary Information section 6), and just a few of the 350 First American individuals sampled exhibited a signal of Paleo-Eskimo ancestry that is comparable to that seen in Na-Dene speakers (Extended Data Fig. 6b). These results suggest that the common ancestor of all Na-Dene branches, now scattered from Arizona and New Mexico to Alaska, experienced Paleo-Eskimo admixture. This scenario is in agreement with evidence from archaeology (see Discussion), and below we further investigate it with explicit demographic modelling based on the rare joint site frequency spectrum.

### No evidence for population turnover in the Aleutian Islands around 1,000 calBP

Morphological disparities between human remains in the Aleutian Islands dated before and after around 1,000 calBP, the time of the Thule expansion, were suggested by Hrdlička as reflecting a population turnover^33^. Archaeological evidence also suggests dramatic material culture changes around this time including burial practices and other cultural expressions^34^, and these distinct cultures were termed Paleo- and Neo-Aleut. Mitochondrial DNA analysis provided some evidence for population turnover via an increase in the frequency of mitochondrial DNA haplogroup D2a1a at the expense of A2a after around 1,0 calBP^35^. However, in the genome-wide ancient DNA that we report here, including 4 samples labeled as Paleo-Aleuts and 7 samples labeled as Neo-Aleuts, we find no evidence for genetic differences among the two groups. This is evident from PCA and *ADMIXTURE* analyses including 2 Paleo-Aleuts and 4 Neo-Aleuts with the highest number of genotyped sites (Fig. 2, Extended Data Fig. 1), in allele frequency differentiation (*F*_ST_ = 0.003 ± 0.002, which is consistent with zero), and in tests for being derived from a homogeneous ancestral population (all statistics of the form D(Outgroup, Test; Neo-Aleut, Paleo-Aleut) which measure whether a Test population shares more alleles with Neo-Aleuts and Paleo-Aleuts are within 3 standard errors of zero). With the *qpWave* method, we also failed to detect additional ancestry in Neo-Aleuts: both groupings were consistent with one stream of ancestry with P-values ranging from 0.089 to 0.395, depending on the outgroups used. We conclude from this that the Aleutian population largely remained continuous during this transition, unlike the transition between Paleo-Eskimos and Neo-Eskimos in much of the mainland (see further discussion in Supplemental Information section 8).

### A Paleo-Eskimo with West Siberian ancestry

The Ust’-Belaya culture of interior Chukotka shows connections with both late Neolithic of interior Siberia (e.g., Bel’kachi, Ymyakhtakh) as well as with Paleo-Eskimo cultures in the Bering Strait region^36,37^. We dated a single burial at the Ust’-Belaya site at the confluence of the Belaya and Anadyr Rivers, from which we also generated genome-wide data (Supplementary Information section 1), and obtained a date of 4,410 – 4,100 calBP (Supplementary Information section 2). Our targeted enrichment approach generated pseudo-haploid genotypes at 832,452 sites across this individual’s genome (Table 1). The position of this sample in the space of two principal components (PC1 and PC2) suggests that it might have ancestry from both Paleo-Eskimos and western Siberian lineages (Fig. 2). Indeed, *qpAdm* analysis demonstrates that the Ust’-Belaya individual can be modelled as descended from Paleo-Eskimos (represented by Saqqaq) and West Siberians (represented by Kets or other groups), with Siberian admixture proportions ranging from ~20 to ~50%, depending on source populations and datasets (Extended Data Fig. 9). Models with East Siberians instead of West Siberians (Extended Data Fig. 9) and/or Chukotko-Kamchatkan speakers instead of Paleo-Eskimos (data not shown) were often inconsistent with two sources of ancestry, or demonstrated negative admixture proportions or very wide error intervals. In line with these findings, the Ust’-Belaya individual is closely related to both Saqqaq and Kets, a West Siberian population, according to *f4*-statistics (Ust’-Belaya, Yoruba; Saqqaq, any other population) and (Ket, Yoruba; Ust’-Belaya, any other population) (Supplementary Table 5). Moreover, this individual has a high level of Mal’ta (ancient North Eurasian, ANE) ancestry according to *f4*-statistics (Mal'ta, Yoruba; Ust’-Belaya, Siberian population) (Supplementary Table 5): an expected result given that West Siberians have substantial ANE ancestry^38^. In summary, striking parallels in archaeological and genetic results suggest that admixture between proto-Paleo-Eskimos and Siberian lineages in Chukotka took place not long after they diverged (see the next section), indicating that cultural contact between these groups at this time almost certainly occurred as well. This result has implications for archaeology and historical linguistics, as discussed below.

Two later individuals from Old Bering Sea culture burials at Uelen, Chukotka, dated at 1,970 – 1,590 calBP and 1,180 – 830 calBP (Supplementary Information section 2), were also subjected to targeted enrichment and sequenced, producing pseudo-haploid genotypes at 608,585 or 797,816 sites (Table 1). Their genetic signature was typical for Neo-Eskimos according to all analyses (see, for example, Figs. 2 and 3). Thus, the older individual represents the earliest Neo-Eskimo for which genetic data were ever reported.

### Demographic modelling

To further interpret our findings, we built an explicit demographic model for the populations analyzed here. We used *rarecoal*^31^ to estimate split times and population sizes, as well as admixture events, in a population tree connecting Europeans, Southeast Asians, Siberians, Chukotko-Kamchatkan, Eskimo-Aleut, and Northern Athabaskan speakers, and southern First Americans. Sample sizes and additional details are provided in Supplementary Information section 9. The model was derived in an iterative way: we started off with fitting a model to three populations only (Europeans, Southeast Asians, and First Americans), and then added one population at a time, re-estimating all previous and new parameters (see details in Supplementary Information section 9). Admixture edges were added when the model fit showed significant deviations for particular allele sharing statistics. We explicitly corrected for Post-Columbian admixture in Chukotko-Kamchatkan and Eskimo-Aleut speakers by adding admixture edges from Europeans into these groups, with a fixed time at 200 calBP (not shown in Fig. 5a). The maximum likelihood parameter estimates for this final model are shown in Supplementary Information section 9.

**Figure 5.**
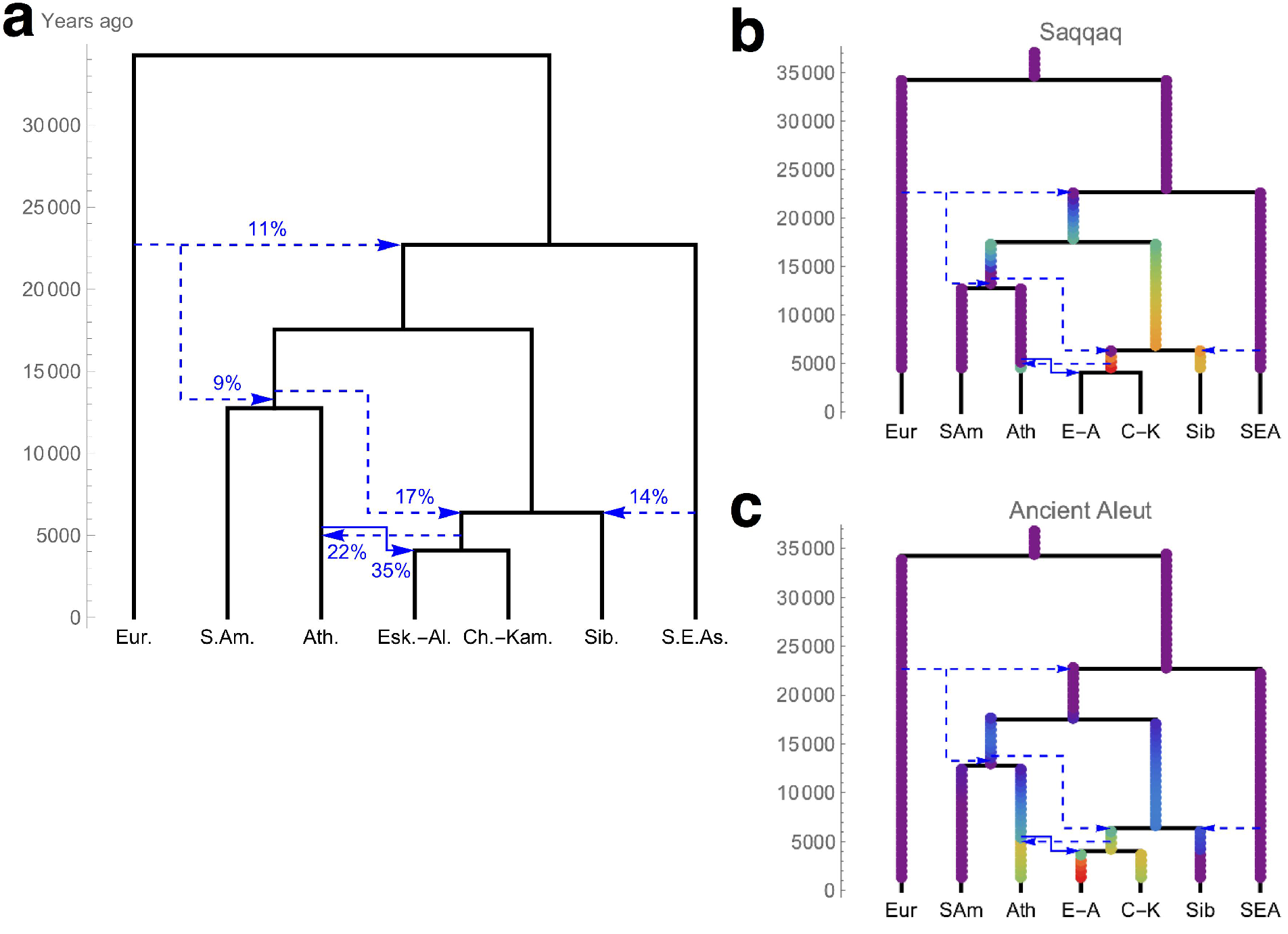
**a**, A demographic model based on 106 individuals from 7 meta-populations, estimated using *rarecoal*, which is a fit to the data. Dashed arrows indicate salient admixture events. For a figure showing all admixture events and for a complete list of parameter estimates see Supplementary Information section 9. Admixture graphs with the same topology are presented in Extended Data Fig. 10 and in Supplementary Information section 9. In the case of European admixture in the Siberian/American and American clades, the arrows indicate a ghost population that split off the European branch around calBP and most likely corresponds to Ancient North Eurasians^39^. A similar ghost population is modelled splitting from the ancestors of Athabaskans and admixing into the branch representing Eskimo-Aleut speakers. We also added admixture edges at 200 calBP from Europeans into some extant groups (Eskimo-Aleut and Chukotko-Kamchatkan speakers), modelling Post-Columbian admixture. These are not shown for clarity. A more ancient European admixture event in the Siberian clade dated at ~4,000 calBP (Supplementary Information section 9) is not shown either. **b** and **c**, Most likely branching points for the Saqqaq and ancient Aleut sample for which we generated complete genome data. Colored points indicate relative log likelihood with respect to the best fitting model. Only branching prior to the radiocarbon dates of the samples was allowed.

Our final model suggests that Arctic populations on both sides of the Bering strait, i.e. Chukotko-Kamchatkan and Eskimo-Aleut speakers, form a clade that separated ~6,300 calBP from the ancestors of present-day Siberians further to the west. The Arctic clade inherited an additional 18% ancestry from the Asian lineage ancestral to Native Americans. We did not attempt to include the ancient Mal'ta genome^39^, which would be a representative of ANE, since *rarecoal* requires high-quality genomes for modelling phylogenies. However, the admixture edge from a European sister group into the ancestor of all American, Siberian, and Arctic groups, and a later admixture exclusive to the Native American lineage (in line with the admixture graphs in Extended Data Fig. 10, see below) most likely reflect ANE admixture.

At 4,000 calBP we infer a split between the ancestors of the Chukotko-Kamchatkan and Eskimo-Aleut speakers, with the latter inheriting a substantial proportion of their ancestry (33%) from an early mixture with a group distantly related to Athabaskan speakers. Finally, we infer that Northern Athabaskans derive 21% of their ancestry from the common ancestor of Chukotko-Kamchatkan and Eskimo-Aleut speakers, likely related to Paleo-Eskimos (see below). We also tested alternative models without this last admixture edge, and with admixture from Eskimo-Aleut or Chukotko-Kamchatkan speakers into Athabaskans, but found substantially poorer fits (Supplementary Information section 9). We note that all time and population size estimates depend linearly on estimates of the human mutation rate (here taken as 0.44 × 10^−9^ per year^40,41^), which has substantial uncertainty. While this model was derived by fitting rare variants of allele frequency up to 1.8%, we also tested whether it fits F-statistics computed for common variants. Specifically, we used *qpGraph*^42^ to test various topologies including the final one derived with *rarecoal* (Extended Data Fig. 10, Supplementary Information section 9), and found that it was indeed consistent with all possible F-statistics in all combinations of meta-populations (the worst Z-score was 0.883).

With a robust maximum likelihood model connecting 7 extant groups, we included two ancient genomes in the modelled tree, correcting for missing genotype calls due to limited coverage in these individuals. We find that the Saqqaq genome^7^ most likely branches off the tree very close to the admixture edge into Northern Athabaskan speakers (Fig. 5b). This positioning of Saqqaq at the ancestral branch of the Arctic clade prior to the Chukotko-Kamchatkan/Eskimo-Aleut split suggest that: 1) Paleo-Eskimos are closely related (but not identical) to the founding population of Neo-Eskimos, and 2) Paleo-Eskimos contributed substantially to Na-Dene speakers. We also mapped the ancient Aleutian Islander, for which we generated whole genome data, onto our fitted tree. We find that this individual is most closely related to present-day Eskimo-Aleut speakers, as seen by the maximum likelihood split point on the ancestral branch of that population. This position confirms the continuity with present-day Aleuts as seen in the PCA and other analyses.

## Discussion

The new data and analyses presented in this work derive two key results on the genetic legacy of Paleo-Eskimos. First, we show that Paleo-Eskimos were very closely related to the Asian founder lineage that gave rise to Eskimo-Aleut speakers. Second, we show that Paleo-Eskimos contributed substantially to the ancestry of Native Americans speaking Na-Dene languages. These results add significantly to previous studies on these topics. Reich *et al.*^1^ inferred that an unspecific Asian source contributed around 43% of the ancestry of Eskimo-Aleut speakers and around 10% of the ancestry of Chipewyans, a Northern Athabaskan-speaking population. Our analyses show that this Asian source is equivalent to the ancestral population that we here term proto-Paleo-Eskimos. We show that within this lineage, two sub-lineages formed that contributed to almost all Na-Dene speakers and to Neo-Eskimos, respectively. According to a different study^2^, Northern Athabaskan speakers did not receive Paleo-Eskimo admixture, but admixture between Athabaskans and Eskimo-Aleut speakers was proposed. While we observe substantial First American ancestry in Eskimo-Aleut speakers, we find no evidence for gene flow from them into Athabaskans. Instead, we propose that the observed genetic patterns can be explained by Paleo-Eskimo ancestry in Athabaskans, as well as in other Na-Dene-speaking populations. Similarly, in a third study^8^, admixture of unresolved direction between Saqqaq and ancestral Neo-Eskimos was interpreted as most likely reflecting Neo-Eskimo admixture into Paleo-Eskimos. Here we show that substantial proto-Paleo-Eskimo ancestry contributed to the founder lineage of Eskimo-Aleut speakers, and think this explains the observed admixture, as well as the presence of mitochondrial haplogroup D2a in the North Slope Iñupiat^22^.

Our results show that Paleo-Eskimo ancestry is a nearly perfect tracer-dye for speakers of Na-Dene languages including the most divergent linguistically (Tlingit) and the most geographically remote ones (Southern Athabaskans, Fig. 6a). It is plausible that Paleo-Eskimos rather than Neo-Eskimos contributed to Na-Dene populations in light of archaeological evidence. The arrival of Neo-Eskimos (the Birnirk and Thule cultures) into western Alaska is dated to 1,350 – 1,150 calBP^8,43^, but at that point Tlingit had probably already come to occupy their current position in southeastern Alaska^16,44^. It has been hypothesized^45–47^ that the spread of Southern Athabaskan speakers from the Subarctic was triggered by a massive volcanic ash fall 1,100 calBP^48^ (Fig. 6a). If this hypothesis is correct, both Tlingit and Apache would have had little opportunity to mix with newly arriving Neo-Eskimos, which would explain why in our analysis, southern Athabaskan speakers and Tlingit have the Paleo-Eskimo ancestry but not the Neo-Eskimo ancestry. In contrast, Paleo-Eskimo peoples lived alongside Na-Dene ancestors for millennia, providing ample opportunity for genetic interaction^16^. Although archaeological evidence for such interaction across the coastal-interior cultural boundary remains sparse^16,46^, our genetic analyses demonstrate that substantial gene flow from Paleo-Eskimos took place (25-40% in ancient Northern Athabaskans).

**Figure 6.**
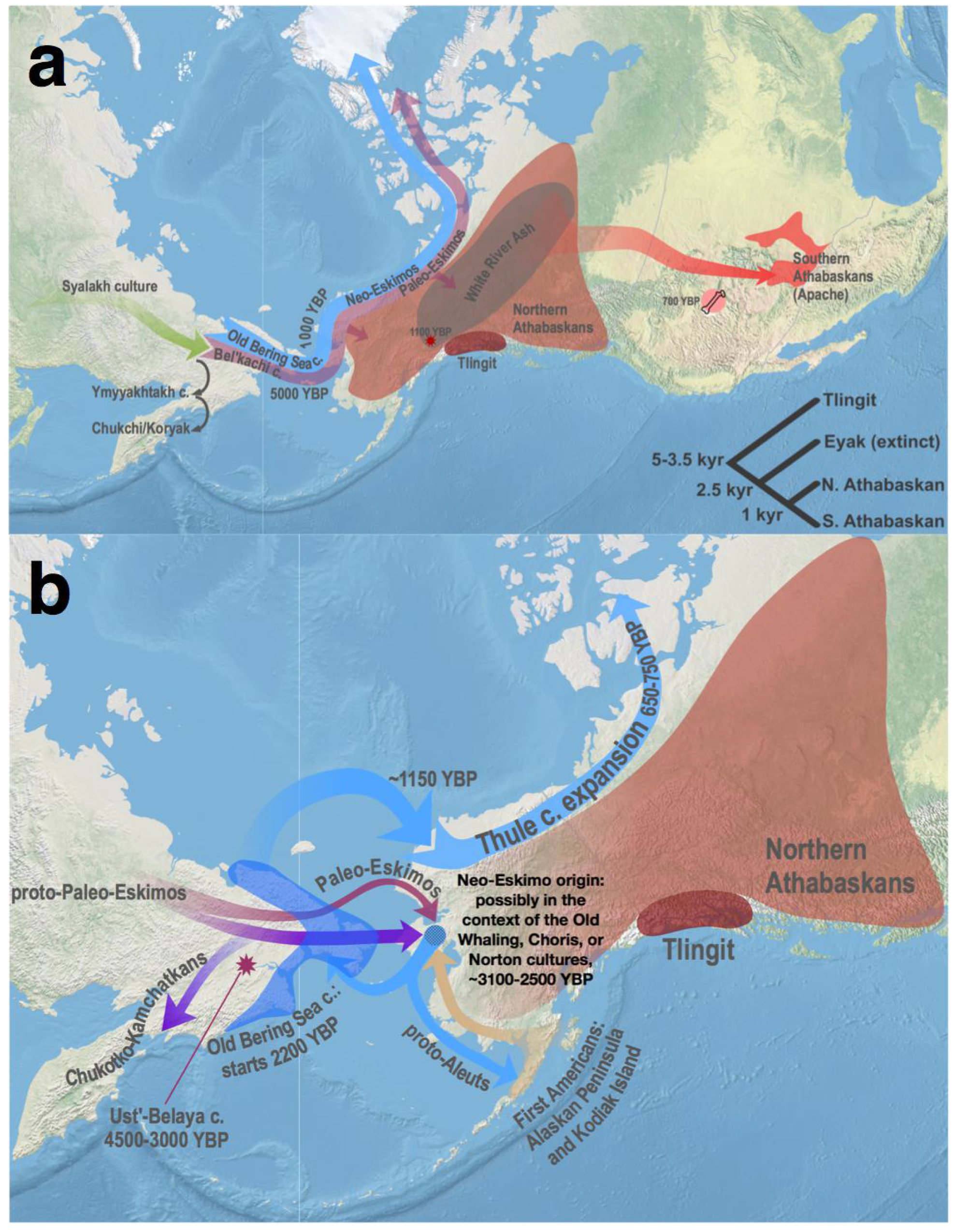
An overview of North American and Chukotkan population history illustrating the history of major Na-Dene groups (**a**) and our model for the emergence and spread of Eskimo-Aleut speakers (**b**). Approximate earliest dates in calBP are indicated for archaeological or ethnic areas and for migrations. Due to space constraints, some migration paths are drawn to indicate just general directions, but not actual routes of population spread. **a**, The Paleo-Eskimo/Na-Dene gene flow we provisionally mapped across the coastal-interior boundary separating the ASTt and Northern Archaic cultures in Alaska, where the highest diversity of Na-Dene languages is found (for that reason Alaska was proposed as a homeland of the Na-Dene language family^67^). The gene flow might take place further east along the same boundary. In addition, this panel shows the succession of archaeological cultures on the Siberian side of the Bering Strait, following the split with Paleo-Eskimos and culminating with present-day Chukchi, Itelmens, and Koryaks. A cladogram of the Na-Dene language family in the bottom right-hand corner shows the current consensus view of language relationships and summarizes published linguistic dating results (see further details in Supplementary Information section 10) The Mount Churchill volcanic eruption that deposited the precisely dated White River Ash^48^ and possibly triggered the departure of Apachean ancestors^45–47,68^ is also shown. While early stages of the Apachean southward migration remain undated, their appearance at the Promontory Caves (Utah) has been dated at 700 – 660 calBP^69^. **b**, A model of population history for Eskimo-Aleut speakers combining genetic and archaeological evidence; see Discussion for details. The Ust’-Belaya site in Chukotka is shown with an asterisk.

The time and place of the Eskimo-Aleut founder event remains uncertain. Under our demographic model, divergence of the lineage leading to Eskimo-Aleut speakers was dated at ~3,500 calBP, and involved gene flow from a northern First American population distantly related to Athabaskans (Fig. 5). There is no clear archaeological evidence for a First American back-migration to Chukotka^16,49^, so this admixture event may have occurred in North America. A parsimonious explanation is that the Asian ancestral population contributing to Eskimo-Aleut speakers may have remained in Chukotka after splitting from the Paleo-Eskimo lineage *sensu stricto*, and that members of this lineage later separated from the ancestors of Chukotko-Kamchatkan speakers and crossed the Bering Strait (Fig. 6b). In turn, the First American ancestral lineage that contributed to Eskimo-Aleut speakers was likely located in southwestern Alaska since the Alaskan Peninsula and Kodiak Island have long been suggested as a source of influences shaping the Neo-Eskimo material culture^37,50^. The earliest maritime adaptations in Beringia and America are encountered in this region associated with the Ocean Bay tradition (~6,800 – 4,500 calBP)^51,52^. Early Paleo-Eskimo people used marine resources on a seasonal basis only, depended for the most part on hunting caribou and muskox, and lacked sophisticated hunting gear that allowed the later Inuit to become specialized in whaling^43^. It is conceivable that a transfer of cultural traits and gene flow happened simultaneously.

Where did the First American and Paleo-Eskimo-related source populations meet? A succession of western Alaskan cultures, namely the Old Whaling, Choris, Norton, and Ipiutak (with the earliest dates around 3,100, 2,700, 2,500, and 1,700 calBP, respectively), combined cultural influences from earlier local Paleo-Eskimo sources as well as sources in Chukotka and southwestern Alaska^37,53^. Parallels between these cultures and subsequent Neo-Eskimos are notable^37^. The Old Bering Sea culture, the earliest culture assigned archaeologically and genetically to Neo-Eskimos^8^, has been dated to around 2,200 calBP and later^20,54^. An individual from Uelen with the Neo-Eskimo genetic signature was dated in this study at ~1,800 calBP. Considering these dates, we provisionally suggest that the admixture that happened early in the history of the Neo-Eskimos may have occurred in the context of the Old Whaling, Choris, or Norton cultures (Fig. 6b), although other scenarios cannot be ruled out without further ancient DNA sampling. It is possible that Paleo-Eskimos *sensu stricto* may have also contributed to some lesser extent to the emergence of Neo-Eskimo peoples (Fig. 6b).

The descendants of proto-Paleo-Eskimos speak widely different languages, belonging to the Chukotko-Kamchatkan, Eskimo-Aleut, and Na-Dene families. Based on lexicostatistical studies of languages surviving in the 20^th^ century, the time depth of the former two families is likely shallow, and the Na-Dene family is probably much older, on the order of 5,000 years (Supplementary Information section 10). Thus, the linguistic affiliation of Paleo-Eskimos is unclear. A Siberian linguistic connection was proposed for the Na-Dene family under the Dene-Yeniseian hypothesis^55,56^. This hypothetical language macrofamily unites Na-Dene languages and Ket, the only surviving remnant of the Yeniseian family, once widespread in South and Central Siberia^57,58^. Perhaps consistent with this hypothesis, one ancient Chukotkan sample from the Ust’-Belaya culture that was first reported in this study shows evidence of ancestry from both Paleo-Eskimos and a western Siberian group related to Kets. This genetic evidence suggests that links across geographic distances such as that between Kets and Paleo-Eskimos may have been possible. Although the Dene-Yeniseian macrofamily is not universally accepted among historical linguists^59,60^, and correlations between linguistic and genetic histories are far from perfect, evidence of a genetic connection between Siberian and Na-Dene populations mediated by Paleo-Eskimos suggests that future research should further explore the genealogical relationships between these language families, either the closest sister-groups^56^ or those within a wider clade^60^.

## Methods

### Ancient DNA sampling, extraction and sequencing

In dedicated clean rooms at Harvard Medical School (the 11 Aleutian Islanders and 3 Tochak McGrath samples), and at University College Dublin (the 3 Chukotkan samples), we prepared powder from human skeletal remains, as described previously^26^. We extracted DNA using the Dabney *et al.*^23^ protocol, and prepared double-stranded barcoded libraries that were treated by UDG to remove characteristic cytosine to thymine damage in ancient DNA using the Rohland *et al.*^24^ protocol. We enriched the libraries for a set of approximately 1.24 million SNPs^25^, and sequenced on a NextSeq instrument using 75 nt paired-end reads, which we merged before mapping to the human reference genome (requiring at least 15 base pairs of overlap) (Supplementary Information section 3). We also carried out shotgun sequencing of one Aleutian Islander individual (Table 1). The work with ancient individuals was conducted only after consultation with local communities and authorities, and after formal permissions were granted. Results have been communicated in person and in writing to descendant communities.

### Sampling present-day populations

Sampling of the Alaskan Iñupiat population (35 individuals) was performed with informed consent as described in Raff *et al.*^22^ (see also Supplementary Information section 1). Saliva samples of four West Siberian ethnic groups (Enets, Kets, Nganasans, Selkups, 58 individuals in total) were collected and DNA extractions were performed as described in Flegontov *etal.*^38^ (see also Supplementary Table 2). Please see ethical approval statements in the respective papers^22,38^.

### Dataset preparation

To analyze rare allele sharing patterns, we composed a set of sequencing data covering Africa, Europe, Southeast Asia, Siberia, and the Americas: 1,207 individuals from 95 populations (Supplementary Table 3). We assembled the dataset using three published sources: the Simons Genome Diversity Project^61^, Raghavan *et al.*^2^, and the 1000 Genomes Project^32^. We used variant calls generated in the respective publications, kept biallelic autosomal SNPs only, and applied the following filtering procedure. We first generated separate masks for the Raghavan *et al.* data and for the SGDP data, based on sites at which at least 90% of all individuals in those data sets have non-missing genotype calls. We then used the overlap of these two masks to generate the final mask for the joint data set. Within this final mask, we treated the few missing genotypes as homozygous reference calls. This was necessary, since in the *rarecoal* analysis we cannot handle missing data, and justified since we are analyzing rare variation, for which a missing genotype is much more likely to be homozygous reference than any other genotype. For the one ancient Aleut individual for which shotgun data was generated, we called variants using a method tailored to rare genetic variants shared with a reference set: at every position in our reference set with allele count below 10, we checked reads overlapping that position. If at least two reads supported the alternative allele, we called a heterozygous genotype. In all other cases, if at least two reads cover a site we called a homozygous reference allele. This method results in a large false negative rate, but relative sharing ratios with reference populations should be relatively unbiased^31^. When analyzing the ancient Aleut together with modern data, we restricted the analysis to regions in which the Aleut sample had non-missing genotypes (i.e. had at least 2x coverage).

Additionally, we assembled two independent SNP datasets: see dataset compositions in Supplementary Table 3 and filtration settings in Supplementary Table 4. Initially, we obtained phased autosomal genotypes for large worldwide collections of Affymetrix HumanOrigins or Illumina SNP array data (Supplementary Table 4), using*Shapeltv.2.20* with default parameters and without a guidance haplotype panel^62^. Then we applied missing rate thresholds for individuals (<50% or <51%) and SNPs (<5%) using *PUNK v.1.90b3.36*^63^. For *ADMIXTURE*, *PCA*, and *qpWave/qpAdm* analyses, more relaxed missing rate thresholds for individuals were applied, 75% or 70% depending on the dataset (Supplementary Table 4). This allowed us to include relevant ancient samples genotyped using the targeted enrichment approach (Supplementary Table 1). For the *ADMIXTURE* analysis, unlinked SNPs were selected using linkage disequilibrium filtering with *PLINK* (Supplementary Table 4). Ten principal components (PC) were computed using *PLINK* on unlinked SNPs, and weighted Euclidean distances defined as:

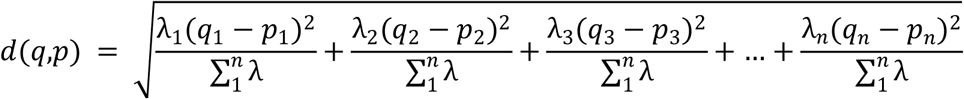

were calculated among individuals within populations (*q*_*n*_ and *p*_*n*_ refer to PCs from 1 to 10 in a population, λ_*n*_ is the corresponding eigenvalue). We removed outliers manually considering the weighted Euclidean distances and results of an unsupervised *ADMIXTURE*^64^ analysis (K=13). Populations having on average >5% of the Siberian ancestral component according to *ADMIXTURE* analysis (Extended Data Fig. 1), e.g. Finns and Russians, were excluded from the European and Southeast Asian meta-populations. In the case of the Illumina SNP array dataset, Na-Dene populations were exempt from PCA outlier removal and from removal of supposed relatives identified by Raghavan *etal.*^2^. This was done to preserve maximal diversity of Na-Dene and to ensure that both Dakelh individuals with sequencing data available would be included; our analysis is designed to be robust to the presence of European admixture. Finally, we selected relevant metapopulations, generating datasets of 489-1161 individuals further analyzed with *ADMIXTURE*^64^, PCA as implemented in *PLINKv.1.90b3.36*^63^, *qpWave/qpAdm*^1,26^, *ChromoPainter v.1* and *fineSTRUCTURE*^29^, *ChromoPainter v.2* and *GLOBETROTTER*^30^ (Supplementary Tables 3 and 4). For the *qpWave/qpAdm* analyses^1,26^, any American individuals with >1% European, African, or Southeast Asian ancestry according to *ADMIXTURE* (Extended Data Fig. 1) were removed, as well as Chukotkan and Kamchatkan individuals with >1% European or African ancestry. Some additional Chipewyan and West Greenlandic Inuit individuals were removed since “cryptic” European ancestry undetectable with *ADMIXTURE* was revealed in them using *D* statistics (Yoruba or Dai, Icelander; Chipewyan individual, Karitiana) and (Yoruba or Dai, Slovak; West Greenlandic Inuit individual, Karitiana). Any individual with any of the two |Z|-scores >3 was removed. The dataset pruning procedure is illustrated on PCA plots presented in Fig. 2, Extended Data Fig. 3, and Supplementary Information section 4.

### *ADMIXTURE* analysis

The *ADMIXTURE* software^64^ implements a model-based Bayesian approach that uses a block-relaxation algorithm in order to compute a matrix of ancestral population fractions in each individual (*Q*) and infer allele frequencies for each ancestral population (*P*). A given dataset is usually modelled using various numbers of ancestral populations (*K*). We ran *ADMIXTURE* on HumanOrigins-based and Illumina-based datasets of unlinked SNPs (Supplementary Table 4) using 10 to 25 and 5 to 20 *K*values, respectively. One hundred analysis iterations were generated with different random seeds. The best run was chosen according to the highest likelihood. An optimal value of *K* was selected using 10-fold crossvalidation.

### Principal component analysis (PCA)

PCA was performed using *PLINK v.1.90b3.36*^63^ with default settings. No pruning of linked SNPs was applied prior to this analysis (Supplementary Table 4), and almost identical results were obtained on pruned datasets.

### Admixture modelling with *qpWave* and *qpAdm*

We used the *qpWave* tool (a part of *AdmixTools*) to infer how many of streams of ancestry relate a set of test populations to a set of outgroups^1^. *qpWave* relies on a matrix of statistics *f*_*4*_(testi, test_*1*_; outgroup_*i*_, outgroup_*x*_). Usually, a few test populations from a certain region and a diverse worldwide set of outgroups (having no recent gene flow from the test region) are co-analyzed^3,26,65^, and a statistical test is performed to determine whether allele frequencies in the test populations can be explained by one, two, or more streams of ancestry derived from the outgroups. If a group of three populations, a triplet, is derived from two ancestry streams according to a *qpWave* test, and any pair of the constituent populations shows the same result, it follows that one of the populations can be modelled as having ancestry from the other two using another tool, *qpAdm*, which makes the implicit assumption that the two populations used as the sources have not undergone admixture^26^.

The following sets of outgroup populations were used for analyses on the HumanOrigins dataset: 1) “9 Asians”, 8 diverse Siberian populations (Nganasan, Tuvinian, Ulchi, Yakut, Even, Ket, Selkup, Tubalar) and a Southeast Asian population (Dai); 2) “19 outgroups” from five broad geographical regions: Mbuti, Taa, Yoruba (Africans), Nganasan, Tuvinian, Ulchi, Yakut (East Siberians), Altaian, Ket, Selkup, Tubalar (West Siberians), Czech, English, French, North Italian (Europeans), Dai, Miao, She, Thai (Southeast Asians); 3) “9 Asians + Koryak”, 8 Siberian populations, Dai, and Koryak, a close outgroup for Americans that should provide higher resolution. The following sets of outgroup populations were used for analyses on the Illumina dataset: 1) “10 Asians”, 9 Siberian populations (Buryat, Dolgan, Evenk, Nganasan, Tuvinian, Yakut, Altaian, Khakas, Selkup) and Dai; 2) “20 outgroups”: Bantu (Kenya), Mandenka, Mbuti, Yoruba (Africans), Buryat, Evenk, Nganasan, Tuvinian, Yakut (East Siberians), Altaian, Khakas, Selkup (West Siberians), Basque, Sardinian, Slovak, Spanish (Europeans), Dai, Lahu, Miao, She (Southeast Asians); 3) ”10 Asians + Koryak”, 9 Siberian populations, Dai, and Koryak. All possible triplets of the form (First American or Na-Dene population; Eskimo-Aleut population; Paleo-Eskimo population) and (First American or Na-Dene pop.; Eskimo-Aleut pop.; Chukotko-Kamchatkan pop.) and quadruplets of the form (First American pop.; Na-Dene pop.; Eskimo-Aleut pop.; Paleo-Eskimo pop.) were tested with *qpWave* on both the HumanOrigins and Illumina SNP array datasets, with or without transition polymorphisms, and using three alternative outgroup sets. Paleo-Eskimos were represented by the Saqqaq or Late Dorset individuals, or by these two individuals combined. For admixture inference with *qpAdm*, all possible triplets of the form (any American, Chukotkan or Kamchatkan pop.; Paleo-Eskimo pop.; Guarani, Karitiana, or Mixe) were considered in the HumanOrigins dataset, and all possible triplets of the form (any American, Chukotkan or Kamchatkan pop.; Paleo-Eskimo pop.; Karitiana, Mixtec, Nisga’a, or Pima) were considered in the Illumina dataset. Paleo-Eskimos were represented by the Saqqaq individual or by the Saqqaq and Late Dorset individuals combined.

### fineSTRUCTURE clustering

We used *fineSTRUCTURE v.2.0.7* with default parameters to analyze the output of *ChromoPainter v.1*^29^. Clustering trees of individuals were generated by *fineSTRUCTURE* based on counts of shared haplotypes^29^, and two independent iterations of the clustering algorithm were performed. The clustering trees and coancestry matrices were visualized using *fineSTRUCTURE GUI v.0.1.0*^29^.

### Haplotype sharing statistics

The Haplotype Sharing Statistic (HSS_*AB*_) is defined as the total genetic length of DNA (in cM) that a given individual *A* shares with individual *B*_*j*_ under the model^29,30^. HSS_*AB*_ was computed in the all vs. all manner by *ChromoPainter v.1*^29^ running with default parameters, and in practice we summed up the length of DNA that individual *A* copied from individual *B*_*j*_ and the length of DNA copied in the opposite direction (from *B*_*j*_ to *A*), i.e. we disregarded the donor/recipient distinction introduced by the *ChromoPainter* software. For each individual *A* (in practice an American individual), HSSab values were averaged across all individuals of a reference population *B* (the Siberian or Arctic meta-population, or the Saqqaq ancient genome^7^), and then normalized by the haplotype sharing statistic HSS_*AC*_ for the European, African, or Siberian outgroup *C*. The resulting statistics HSS_*AB*_/HSS_*AC*_ are referred to as Siberian, Arctic, or Saqqaq relative haplotype sharing, and were visualized for separate individuals. Similar statistics were calculated for Siberian and Arctic individuals using the leave-one-out procedure. Relative HSSs for recently admixed populations, with ancestry from population *A* and population *B*, were calculated in the following way: *a*×HSS_AC_/HSS_AD_ + *b*×HSS_BC_/HSS_BD_, where *a* and *b* are admixture proportions being simulated.

### Dating admixture events using haplotype sharing statistics

We used *GLOBETROTTER*^30^ to infer and date up to two admixture events in the history of Na-Dene populations. To detect subtle signals of admixture between closely related source populations, we followed the ‘regional’ analysis protocol of Hellenthal *etal.*^30^ Using *ChromoPainter v.2*^30^, chromosomes of a target Na-Dene population were ‘painted’ as a mosaic of haplotypes derived from donor populations or meta-populations: the Saqqaq ancient genome, Chukotko-Kamchatkan groups, Eskimo-Aleuts, northern First Americans, southern First Americans, West Siberians, East Siberians, Southeast Asians, and Europeans. Target individuals were considered as haplotype recipients only, while other populations or meta-populations were considered as both donors and recipients. That is different from the *ChromoPainter v.1* approach, where all individuals were considered as donors and recipients of haplotypes at the same time, and only self-copying was forbidden.

Painting samples for the target population and ‘copy vectors’ for other (meta)populations called ‘surrogates’ served as an input of *GLOBETROTTER*, which was run according to section 6 of the instruction manual of May 27, 2016. The following settings were used: no standardizing by a “NULL” individual (null.ind 0); five iterations of admixture date and proportion/source estimation (num.mixing.iterations 5); at each iteration, any surrogates that contributed ≤ 0.1% to the target population were removed (props.cutoff 0.001); the x-axis of coancestry curves spanned the range from 0 to 50 cM (curve.range 1 50), with bins of 0.1 cM (bin.width 0.1). Confidence intervals (95%) for admixture dates were calculated based on 100 bootstrap replicates. Alternatively, when using separate populations as haplotype donors, the setting ‘standardizing by a “NULL” individual’ was turned on to take account for potential bottleneck effects. A generation time of 29 years was used in all dating calculations^2^.

The *GLOBETROTTER* software is able to date no more than two admixture events^30^, and we therefore had to reduce the complexity of original Na-Dene populations that likely experienced more than two major waves of admixture. For that purpose, only a subset of Na-Dene individuals was used for the *GLOBETROTTER* analysis: those with prior evidence of elevated Paleo-Eskimo ancestry (Supplementary Information section 6) and with <10% West Eurasian ancestry estimated with *ADMIXTURE* (Extended Data Fig. 1).

### Rare allele sharing statistics

We define the Rare Allele Sharing Statistic (RASS_*AB*_) as the average number of sites at which an individual *A* shares a derived allele of frequency *k* with an individual from population *B*:

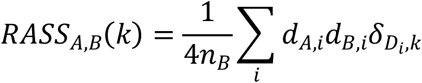

where *n*_*B*_ the number of individuals in population *B*, *d*_*A,i*_ stands for the number of derived alleles at site *i* in individual *A*, and the term *δ*_*x,y*_ equals 0 if the total count of derived alleles in the dataset does not equal *k*, and is 1 otherwise. The sum across all sites *i* is normalized by the product of population sizes multiplied by four to give the average number of shared alleles between two randomly drawn haploid chromosome sets. Instead of counting derived alleles, in practice we counted non-reference alleles, which should not make a difference for low frequencies. To take care of variability in genome coverage across individuals and of dataset-specific SNP calling biases, we calculated normalized (or relative) RASS, dividing RASS_*AB*_ by RASS_*AC*_, where population *C* is a distant outgroup. Standard deviation of RASS_*AB*_ was calculated with a jackknife approach. Specifically, we re-estimated RASS per one million base pairs for drop-one-out data sets excluding an entire chromosome each time. We then used the weighted jackknife method^66^ to estimate sample variances across the drop-one-out data sets. The standard deviation of normalized RASS was calculated using error propagation via partial derivatives:

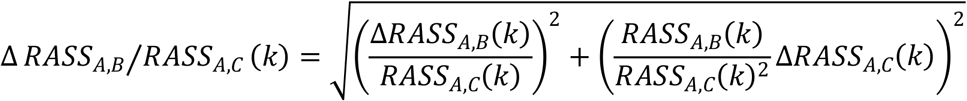

In practice, individual *A* was a present-day or ancient American, population *B* was represented by Siberian or Arctic meta-populations or by the Saqqaq ancient genome^7^, and population *C* - by Africans or Europeans (Supplementary Table 3). The resulting statistics are referred to as relative Siberian, Arctic or Saqqaq allele sharing. Similar statistics were calculated for Siberian and Arctic individuals using the leave-one-out procedure. The same statistics were calculated on a dataset without transition polymorphisms. The ancient Aleut and Saqqaq ancient genomes were not included into the Arctic reference metapopulation. Relative RASS for recent mixtures of individual *A* and individual *B* were calculated in the following way: *a*×RASS_AC_/RASS_AD_ + *b*×RASS_BC_/RASS_BD_, where *a* and *b* are admixture proportions being simulated.

### Demographic modelling

We used the *qpGraph* method^42^ to explore models that are consistent with F statistics and arrived at a final model connecting 8 groups: Mbuti, French, Ami, Mixe, Even, Yupik Naukan, Koryak and Chipewyan (discussed in Supplementary Information section 9). We then used the *rarecoal* program^31^ (https://github.com/stschiff/rarecoal) to derive a timed admixture graph for meta-populations (Fig. 5 and Supplementary Information section 9). We started with a tree connecting Europeans, Southeast Asians, and southern First Americans into a simple tree without admixture, and used “*rarecoal mcmc*” to infer maximum likelihood branch population sizes and split times. We then iteratively added Core Siberians, Chukotko-Kamchatkan, Eskimo-Aleut, and Northern Athabaskan speakers. After each addition, we re-optimized the tree and inspected the fits of the model to the data. When we saw a significant deviation between model and data for a particular pairwise allele sharing probability, we added admixture edges (Supplementary Information section 9). After *rarecoal’s* inference, we rescaled time and population size parameters to years and real effective population size using a mutation rate of 1.25×10^−8^ per site per generation, and a generation time of 29 years^2^. We finally tested whether our final model (Fig. 5a) was consistent also with F statistics using *qpGraph* (Supplementary Information section 9). In order to map the two ancient genomes, Saqqaq and Aleut, we used “*rarecoal find*” to explore a set of possible split points of the ancient lineage on the tree, distributed across all branches and times. Here we restricted the analysis to variants between allele counts 2 and 4. We excluded singletons to reduce impact of false positive genotyping calls^31^.

## Acknowledgements

We gratefully acknowledge the Aleut Corporation, the Aleut/Pribilof Island Association, and the Chaluka Corporation for granting permissions to conduct genetic analyses on the eastern Aleutian remains to help elucidate the population history of the region. We also thank the staff at the Smithsonian Institution’s National Museum of Natural History for facilitating the sample collection. Sample collection and the initial molecular, isotopic and AMS ^14^C dating of the samples described here were funded by National Science Foundation Office of Polar Program grants OPP-9726126, OPP-9974623, and OPP-0327641, by the Natural Sciences and Engineering Research Council of Canada, and the Wenner-Gren Foundation for Anthropological Research (#6364). We are also grateful to the McGrath Native Village Council and MTNT Ltd. for granting permissions to conduct genetic analyses on the Tochak McGrath remains, and to Jamie Clark, who performed biological age estimates on these remains. We thank the research participants in Alaska who donated samples for genome-wide analysis. We are grateful to all researchers that shared their data: Maanasa Raghavan, Simon Rasmussen, Eske Willerslev, and Joan Brenner Coltrain. We also acknowledge valuable advice on American archaeology from Ben A. Potter, John W. Ives and T. Max Friesen. We thank Justin Tackney, Lauren Norman, and Kim TallBear for comments on earlier drafts of this paper. P.F. and E.A. were supported by the Institutional Development Program of the University of Ostrava and by EU structural funding Operational Programme Research and Development for Innovation, project No. CZ.1.05/2.1.00/19.0388. P.C. was supported by the grant no. 0924/2016/ŠaS from the Statutory City of Ostrava and by the grant no. 01211/2016/RRC 'Strengthening international cooperation in science, research and education' from the Moravian-Silesian Region. D.R. was funded by NSF HOMINID grant BCS-1032255, NIH (NIGMS) grant GM100233, and is an Investigator of the Howard Hughes Medical Institute. D.A.B. was supported by a Norman Hackerman Advanced Research Program (NHARP) grant from the Texas Higher Education Coordinating Board (THECB). AMS ^14^C work at Pennsylvania State University was funded by the NSF Archaeometry program award BCS-1460369 to D.J.K.

## Author contributions

S.S., P.F., and D.R. supervised the study. A.M.K., R.A.S., S.V., E.V., D.H.O’R., R.P., and D.R. assembled the collection of archaeological samples. D.A.B., O.F., J.R., M.G.H., and J.K. assembled the sample collection from present-day populations. T.K.H. and D.J.K. were responsible for radiocarbon dating and calibration. N.R., F.C., and D.K. performed laboratory work and supervised ancient DNA sequencing. P.F., N.E.A., P.C., S.M., C.J., T.C.L., I.0., P.S., and S.S., analyzed genetic data. E.J.V. wrote the supplemental section on linguistics. P.F., D.R., and S.S. wrote the manuscript with additional input from all other co-authors.

## Author information

Raw sequence data (bam files) from the 17 newly reported ancient individuals is available from the European Nucleotide Archive. The accession number for the sequence data reported in this paper is (to be provided prior to publication). The genotype data for the Iñupiat are obtained through informed consents that are not consistent with public posting of the data, analyses of phenotypic traits, or commercial use of the data. In order to protect the privacy of participants and ensure that their wishes with respect to data usage are followed, researchers wishing to use data from the Iñupiat samples should contact Geoffrey Hayes (http://ghayes@northwestern.edu) and Deborah Bolnick

(http://deborah.bolnick@austin.utexas.edu), who can then arrange to share the data with researchers who can formally affirm that they will abide by these conditions. The newly reported SNP genotyping data for West Siberians (Enets, Ket, Nganasan, Selkup) is available to researchers who send a signed letter to J.K., P.F., and D.R. containing the following text: “(a) I will not distribute the data outside my collaboration; (b) I will not post the data publicly; (c) I will make no attempt to connect the genetic data to personal identifiers for the samples; (d) I will not use the data for commercial purposes.” The programming code used in this study is available at https://github.com/stschiff/rarecoal. The authors declare no conflicting financial interests. Correspondence and requests for materials should be addressed to S.S. (http://schiffels@shh.mpg.de), P.F. (http://pavel.flegontov@osu.cz), and D.R. (http://reich@genetics.med.harvard.edu).

